# AntiRef: reference clusters of human antibody sequences

**DOI:** 10.1101/2022.12.30.522338

**Authors:** Bryan Briney

## Abstract

**Motivation:** Biases in the human antibody repertoire result in publicly available antibody sequence datasets containing many duplicate or highly similar sequences. These redundant sequences are a barrier to rapid similarity searches and reduce the efficiency with which these datasets can be used to train statistical or machine learning models of human antibodies. Identity-based clustering provides a solution; however, the extremely large size of available antibody repertoire datasets makes such clustering operations computationally intensive and potentially out of reach for many scientists and researchers who would benefit from such data.

**Results:** AntiRef (Antibody Reference Clusters), which is modeled after UniRef, provides clustered datasets of filtered human antibody sequences. Due to the modular nature of recombined antibody genes, the clustering thresholds used by UniRef to cluster general protein sequences (100, 90, and 50 percent identity) are suboptimal for antibody clustering. Starting from a dataset of ∼451 million full-length, productive human antibody sequences from the Observed Antibody Space (**OAS**) repository, AntiRef provides antibody sequence datasets clustered at a range of identity thresholds better suited to antibody sequences. AntiRef90, which uses the least stringent clustering threshold, is roughly one-third the size of the input dataset and less than half the size of the non-redundant AntiRef100.

**Availability:** AntiRef is freely available via Zenodo (zenodo.org/record/7474336) under the CC-BY 4.0 license, which matches the licensing of OAS datasets. All code used to download and filter sequences, generate AntiRef clusters, and create the figures used in this publication is freely available under the MIT license via GitHub (github.com/briney/antiref). We anticipate AntiRef updates will be released bi-annually, with the option for supplementary out-of-band updates when large or particularly interesting datasets are made available. The AntiRef versioning scheme (current version: v2022.12.14) refers to the date on which sequences were retrieved from OAS.

## Introduction

The massive diversity of the human antibody (**Ab**) repertoire is produced initially by somatic recombination of germline gene segments [1]. Ab heavy chains (**HCs**) are assembled from variable (**V**), diversity (**D**) and joining (**J**) gene segments, with D to J recombination followed by V to DJ recombination (***Figure 1***). Light chain (**LC**) recombination proceeds similarly, although without D gene segments. Considering both heavy and light chains, it is estimated that the VDJ recombination process can generate as many as 10^18^ unique Abs [2]. For perspective, this surpasses the combined number of unique proteins encoded by all the genomes of all species on earth by many orders of magnitude [3]. Upon exposure to an infectious agent, pathogen-specific Abs are affinity matured in germinal centers [4–6]. Affinity maturation is an iterative process consisting of multiple rounds of clonal expansion, somatic hypermutation (**SHM**) and antigen-driven selection capable of improving Ab affinity by several orders of magnitude [7]. As a result of the affinity maturation process, antigenic stimulation of a single naive B cell can produce a clonal lineage of many B cells, each expressing an Ab that is related to the parental Ab but which has accumulated a unique set of mutations. Following pathogen clearance, a subset of B cells encoding affinity matured Abs are retained as an *immune memory* of the pathogen encounter [8,9]. The formation of this immune memory allows the humoral immune system to rapidly respond to subsequent exposures and is the primary mechanism of protection for virtually all available vaccines. In effect, an individual’s unique collection of affinity matured Ab genes constitutes a complete molecular record of all previous pathogen encounters and encodes all the functional information necessary to prevent infection by pathogens against which an individual is immune.

**Figure 1.**
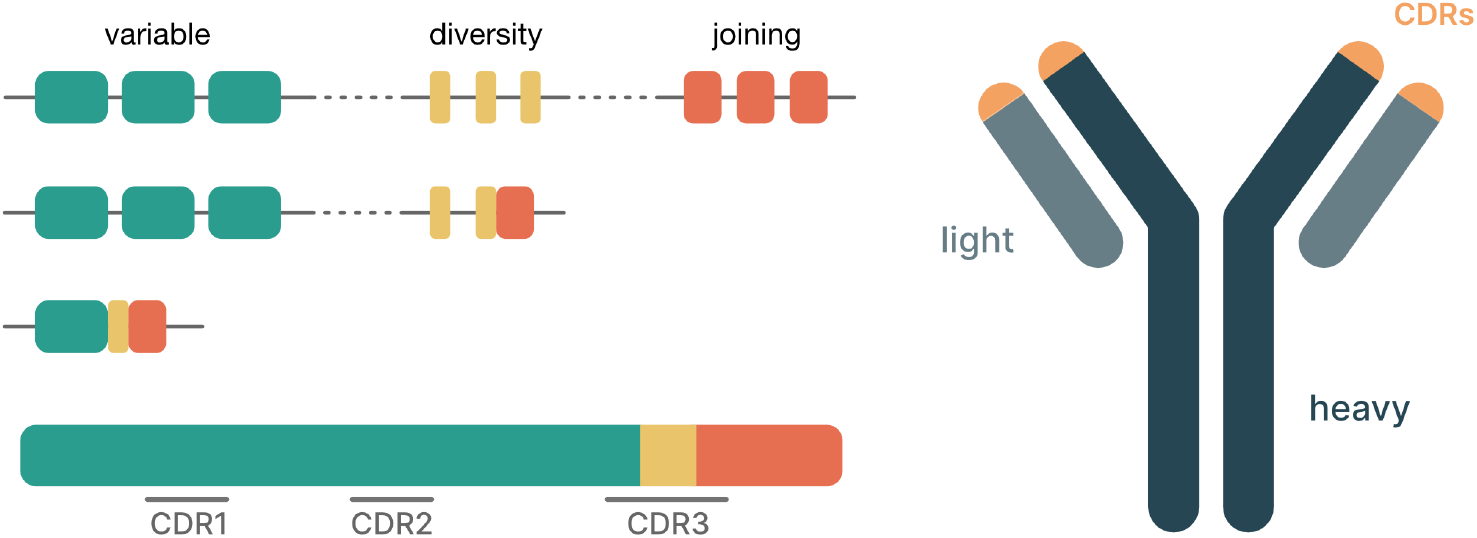
Generation of Ab diversity. The heavy chain VDJ recombination process is shown on the left, with D-J recombination preceding V-DJ recombination. At bottom left is a recombined HC, with approximate CDR locations indicated. On the right is a cartoon representation of an Ab protein, with approximate CDR locations indicated.

High-throughput genetic analysis of antibody repertoires, which became technically possible with read-length enhancements on the 454 GS-FLX in 2009 [10], unlocked fundamental breakthroughs in our understanding of human humoral immunity. Over the ensuing decade, next-generation sequencing (**NGS**) technology has continued to improve [11,12], enabling the first ultra-deep analyses of the human antibody repertoire using billions of sequencing reads [2,13]. These increasingly large antibody repertoire datasets have been reused in a variety of ways, including mining naturally-occurring repertoires for homologs to therapeutic antibodies [14], identification of convergent antibody responses to infection or vaccination [15–18], and identifying vaccine-targetable precursors of exceptionally broad and potent antiviral antibodies [19–23]. One of the most exciting areas of emerging research is the development of sophisticated machine learning models of antibodies and antibody repertoires. Following in the footsteps of large language models for text [24,25] and general protein sequences [26], antibody-specific language models have begun learning features unique to antibodies [27–29] and can be fine-tuned to perform downstream tasks such as structure and paratope prediction [27,30].

These studies are hampered, however, by the scale of available antibody sequence data and the lack of standardized datasets of substantially reduced size that maintain an accurate representation of overall diversity. The availability of such datasets is vitally important; one particularly relevant recent example is the use of UniRef datasets [31,32] to train the state-of-the-art protein language model ESM-2 [26]. We lack a “UniRef for antibodies” in part because its creation is sufficiently computationally intensive to be infeasible for many who would nevertheless benefit from such a dataset. Here, we present AntiRef, a UniRef-inspired, standardized dataset of clustered human antibody sequences.

## System and Methods

### Source data

The antibody sequences used to construct AntiRef were downloaded from the Observed Antibody Space (**OAS**) repository [33,34]. Sequences were filtered using OAS’s query tool prior using the following criteria:

- Species: **Human**
- BSource: **PBMC**
- Disease: **None**
- Vaccine: **None**

All sequences that met these criteria were retrieved using the download files generated by OAS. This resulted in a total of 903,519,744 sequences: 631,028,215 heavy chains and 272,491,529 light chains.

### Sequence filtering

After download, sequences were further filtered using OAS annotation information to retain only sequences with:

- a complete V(D)J region
- no V-gene frameshift insertions or deletions
- in-frame V and J genes
- no stop codons
- no ambiguous amino acids

Following these post-download filtering steps, the dataset contained 260,373,862 heavy chains and 190,684,852 light chains, for a total of 451,058,708 human antibody sequences. Each sequence was given a unique identifier using Python’s built-in `uuid.uuid4()` function, and the complete amino acid sequence of the V(D)J region was assembled from the translated sequence regions in OAS’s annotated output.

### Sequence clustering

Filtered heavy and light chain sequences were pooled and iteratively clustered using the `linclust` function in MMseqs2 [35]. Clustering operations were ordered such that the identity threshold decreased with each round. Following each clustering iteration, representative sequences for each cluster were retrieved and used as input for the subsequent round. This process mirrors the strategy used by UniRef and ensures continuity of sequence and cluster names across all AntiRef datasets [31].

### Incremental AntiRef updates

To create updated versions of AntiRef without requiring a complete re-clustering, new sequences (those which have become available on OAS after the download date of the current AntiRef version) will be downloaded, filtered, and uniquely named as described above. New sequences will be added to the set of sequences comprising the most recent AntiRef100 dataset, and an updated version of AntiRef100 will be generated using the `clusterupdate` function in MMseqs2. Iterative clustering operations will then proceed as described above to generate the remaining AntiRef datasets.

## Results and Discussion

### Database size reduction

A primary motivation for creating AntiRef was to facilitate faster similarity searches across the vast number of publicly available antibody sequences from healthy human donors. By collapsing redundant or highly similar sequences, much like the approach used by UniRef, we can construct smaller datasets that are more easily searchable while still representative of the overall available antibody sequence diversity. The size of each AntiRef dataset, in number of sequences and as a fraction of the total input data, is shown in ***Table 1***. Notably, AntiRef100, which contains all unique sequences from the filtered OAS data, compressed the filtered input dataset by over 25%. AntiRef90, which uses the least stringent clustering threshold of all AntiRef datasets, is 65% smaller than the filtered input dataset. When compared to UniRef, the reduction in size of AntiRef90 (65%) is similar to the compression achieved by UniRef50 (70%), indicating that the antibody-specific clustering thresholds used for AntiRef produce results proportionate to the general protein thresholds used by UniRef.

**Table 1.** AntiRef dataset sizes.

| dataset | clusters | % of input | % of AntiRef100 |
| --- | --- | --- | --- |
| AntiRef100 | 335,140,854 | 74.3% | 100.0% |
| AntiRef99 | 316,067,031 | 70.1% | 94.3% |
| AntiRef98 | 256,587,102 | 56.9% | 76.6% |
| AntiRef96 | 210,600,423 | 46.7% | 62.8% |
| AntiRef94 | 184,479,216 | 40.9% | 55.0% |
| AntiRef92 | 170,094,948 | 37.7% | 50.8% |
| AntiRef90 | 159,871,557 | 35.4% | 47.7% |

### Distribution of cluster sizes

Cluster size frequencies in UniRef datasets follow a power law distribution, meaning the UniRef clustering approach effectively increases dataset diversity by collapsing highly similar sequences that are (in some cases, massively) over-represented. As a result, the general protein language model ESM-2 demonstrated marked improvement when trained with UniRef clusters rather than raw sequence data (Lin *et al*., 2022). Cluster size frequencies in each AntiRef clustering dataset showed a power law distribution (***Figure 2***), providing additional evidence that the benefits of using UniRef datasets in the general protein space can be replicated when using AntiRef datasets in antibody-specific contexts.

**Figure 2.**
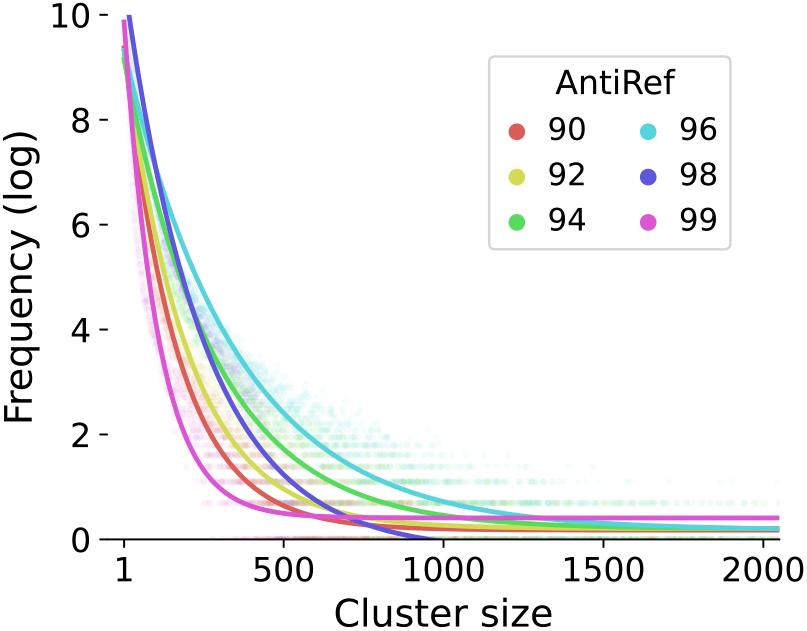
AntiRef cluster size frequencies follow a power law distribution. Cluster sizes were computed for each AntiRef dataset, and the frequency of each cluster size is plotted. The best fit line of the cluster size frequency distribution was separately computed for each AntiRef dataset in Python using `scipy.optimize.best_fit()`.

### Database content and availability

The complete AntiRef package comprises several components:

- *AntiRef annotations:* a single CSV file, containing all filtered and renamed antibody sequences used to construct AntiRef clusters. The CSV also includes the complete set of annotations provided by OAS.
- *AntiRef cluster manifest:* a single CSV file, containing the sequence and cluster IDs for each AntiRef dataset. Because we use a nested clustering approach and name each cluster after its representative sequence, the sequences comprising any cluster from any AntiRef dataset can be easily retrieved.
- *AntiRef data files:* a single file for each AntiRef dataset containing FASTA-formatted amino acid sequences.
- *AntiRef download files:* a pair of text files (one each for heavy and light chains) containing all OAS download commands needed to retrieve all sequence data used to build AntiRef.

All AntiRef datasets are available via Zenodo [37–44] under the CC-BY 4.0 license, which matches the license under which OAS data are released. All code used to generate AntiRef is available via GitHub (github.com/briney/antiref) under the MIT license.

## Conclusions

Inspired by the usefulness of UniRef databases for various computational analyses of general proteins, AntiRef has been created to perform a similar task for antibody sequences. A series of AntiRef datasets have been generated using a nested clustering approach using identity thresholds selected for their appropriateness for antibody sequence data. The regular update cycle will ensure that AntiRef datasets are representative of currently available antibody sequence data, and the provision of all code needed to reproduce AntiRef allows future work extending the AntiRef concept to additional species or to T cell receptor repertoires.

## Acknowledgements

This work was funded by the National Institutes of Health awards R35-GM133682, U19-AI135995, R01-AI171438, and UM1-AI144462. This work would not have been possible without the Observed Antibody Space repository, which facilitates simple and rapid retrieval of large numbers of antibody sequence datasets.

## Notes

https://zenodo.org/record/7474336

https://github.com/briney/antiref

### Competing Interest Statement

The authors have declared no competing interest.

